# Nucleic Acid Capture from Human Blood Plasma Uncovers G-Quadruplex Structures

**DOI:** 10.64898/2026.09.05.747829

**Authors:** Martin Gajarský, Olivia van Ray, Thomas Akkermann, Pascal Hunold, Anne Cucchiarini, Jean-Louis Mergny, Lukáš Trantírek, Robert Hänsel-Hertsch

## Abstract

Ultrashort (US) cell-free DNA (cfDNA) is a population of approximately 50-nucleotide single-stranded DNA molecules in human plasma.^1–3^ Although it holds biological and diagnostic potential, US cfDNA escapes detection by conventional double-stranded library preparation methods.^1–3^ Independent studies have linked US cfDNA to regulatory genomic regions and to sequences predicted to form noncanonical structures, prompting the hypothesis that higher-order DNA structure contributes to its molecular properties. This interpretation, however, has so far rested on computational prediction rather than experimental evidence. Here we test this hypothesis using computational, biophysical and biochemical approaches. *In silico* size-selected US cfDNA from 20 healthy donors was selectively enriched at putative quadruplex sequences (PQS) that overlap both experimentally observed quadruplex sequences and accessible chromatin of blood cells. Synthetic oligonucleotides corresponding to the most enriched of these loci adopted predominantly parallel G-quadruplex (G4) structures, as revealed by circular dichroism. Endogenous nucleic acids captured directly from pooled plasma by poly(A)-tailing and immobilization, without extraction, denaturation or annealing at any step, displayed folded G4 structures. Two orthogonal probes detected these structures: the BG4 antibody and the fluorogenic ligand N-methyl mesoporphyrin IX. Reciprocal competition with a third, chemically unrelated G4 ligand, pyridostatin, confirmed the signal. Together, these experiments provide direct experimental evidence for the presence of folded G4 structures in human blood plasma.

## Introduction

Cell-free DNA (cfDNA) is a central component of modern liquid biopsy, giving minimally invasive access to genomic and epigenomic information associated with physiological and pathological processes. Beyond mutation detection, fragmentomic analyses have shown that fragment size, genomic distribution, end motifs, nucleosome positioning and methylation together encode biological information extending well beyond DNA sequence alone.^4–6^

A major advance in this field was the independent discovery of ultrashort (US) cfDNA, a previously unrecognized population of approximately 50-nucleotide single-stranded DNA molecules that escapes conventional double-stranded library preparation^1–3^. Despite differences in sample preparation and sequencing workflow, these studies agreed that US cfDNA is a molecular population distinct from conventional mononucleosomal cfDNA, with potential utility for liquid biopsy.

Subsequent work characterized this population in more detail. Genome-wide fragmentomic analyses showed enrichment of US cfDNA corresponding to promoter sequences and other regulatory elements, identified characteristic fragmentation and end-motif signatures, and highlighted its potential for cancer detection^2,7^. Methylome analysis and strand-specific sequencing further revealed distinct methylation landscapes, tissue-of-origin information and transcription-associated strand asymmetry^8,9^. Together these studies establish ultrashort plasma DNA as a biologically distinct component of the circulating nucleic acid pool.

A recurrent observation across these studies is the association of US cfDNA with genomic regions capable of forming noncanonical DNA structures. Hisano *et al*. predominantly identified complementary C-rich molecules, reflecting their single-stranded library preparation, whereas Hudecova *et al*. and subsequent studies reported enrichment of G-rich, chromatin-accessible putative quadruplex sequences (PQS) predicted *in silico*; both models nevertheless converge on promoter-associated regions^1,2,7^. G-quadruplex (G4) formation has consequently become an attractive structural model for the biological properties of US cfDNA.

G4s are four-stranded nucleic acid secondary structures built from stacked guanine tetrads, each tetrad held together by Hoogsteen hydrogen bonding and the stack stabilized by a monovalent cation, most effectively potassium, coordinated between successive tetrads. Sequencing of genomic DNA under quadruplex-promoting conditions identified several hundred thousand observed quadruplex sequences (OQS) in the human genome^10^. Antibody-based mapping in chromatin subsequently showed that only a minority of these loci are folded in cells, and that the folded fraction concentrates in nucleosome-depleted, transcriptionally active promoters and enhancers^11,12^. G4 folding is therefore conditional rather than obligate, depending on sequence, strand accessibility and ionic environment.

G4 formation is not confined to the intracellular compartment. Extracellular DNA in bacterial biofilms adopts G4 conformations that resist DNase I and contribute to the stability of the biofilm matrix^13^. In mammalian systems, neutrophil extracellular traps, an established source of circulating cfDNA, contain BG4-reactive G4s that survive nuclease challenge and form catalytically active DNAzymes in complex with heme^14^. Extracellular nucleic acids can therefore maintain quadruplex folding outside the cell, and such folding can confer nuclease resistance: a plausible mechanism by which structured fragments might be selectively retained in a nuclease-rich compartment such as plasma.

Whether nucleic acids circulating in human blood plasma actually adopt folded G4 structures has nevertheless not been established by direct, controlled measurement. Fragmentomic sequencing defines sequence composition, genomic localization and fragmentation patterns but not molecular conformation, and computational prediction of G4-forming motifs establishes structural potential rather than the presence of folded structures.

Here we test the proposed G-quadruplex model of US cfDNA. We first asked whether genomic loci containing chromatin-accessible PQS are particularly enriched in the US cfDNA of healthy donors and examined the structure-forming properties of the most enriched sequences by circular dichroism (CD). We then developed a method to capture endogenous nucleic acids directly from plasma by poly(A)-tailing and immobilization, and to visualize them using chemically unrelated G4 probes. Together these experiments provide direct experimental evidence for folded G-quadruplex structures in human blood plasma.

## Results

### US cfDNA is enriched at putative quadruplex sequences that fold into G4 structures in vitro

Fragmentomic analyses have consistently associated US cfDNA with G4-forming sequences predicted *in silico*^1,7^, but sequence composition alone does not establish whether representative US cfDNA sequences can adopt folded conformations. To identify PQS representative of US cfDNA, we ranked biologically relevant PQS loci by their median CPM-normalized US cfDNA coverage across 26 libraries from 20 healthy donors. Because genomic G4 formation occurs predominantly in accessible chromatin^11^, we reasoned that translocation of these sequences to plasma may reflect cell death, during which they are excised from the open chromatin of dying cells by endogenous DNases. Paired-end aligned whole-genome sequencing data from healthy donors were size-selected *in silico* to fragments of 30–70 bp, and coverage was computed across annotated PQS regions meeting three independent criteria: overlap with an experimentally observed quadruplex sequence (OQS) from G4-seq,^10^ overlap with open chromatin of blood cell types and location within a promoter interval (transcription start site ± 1 kb) (hereafter PQS/OQS/OC). Of 8,316 qualifying intervals, the 50 receiving the highest US cfDNA coverage are shown in Fig. 1A, defining a subset of PQS/OQS/OC loci preferentially enriched in US cfDNA.

**Fig. 1:**
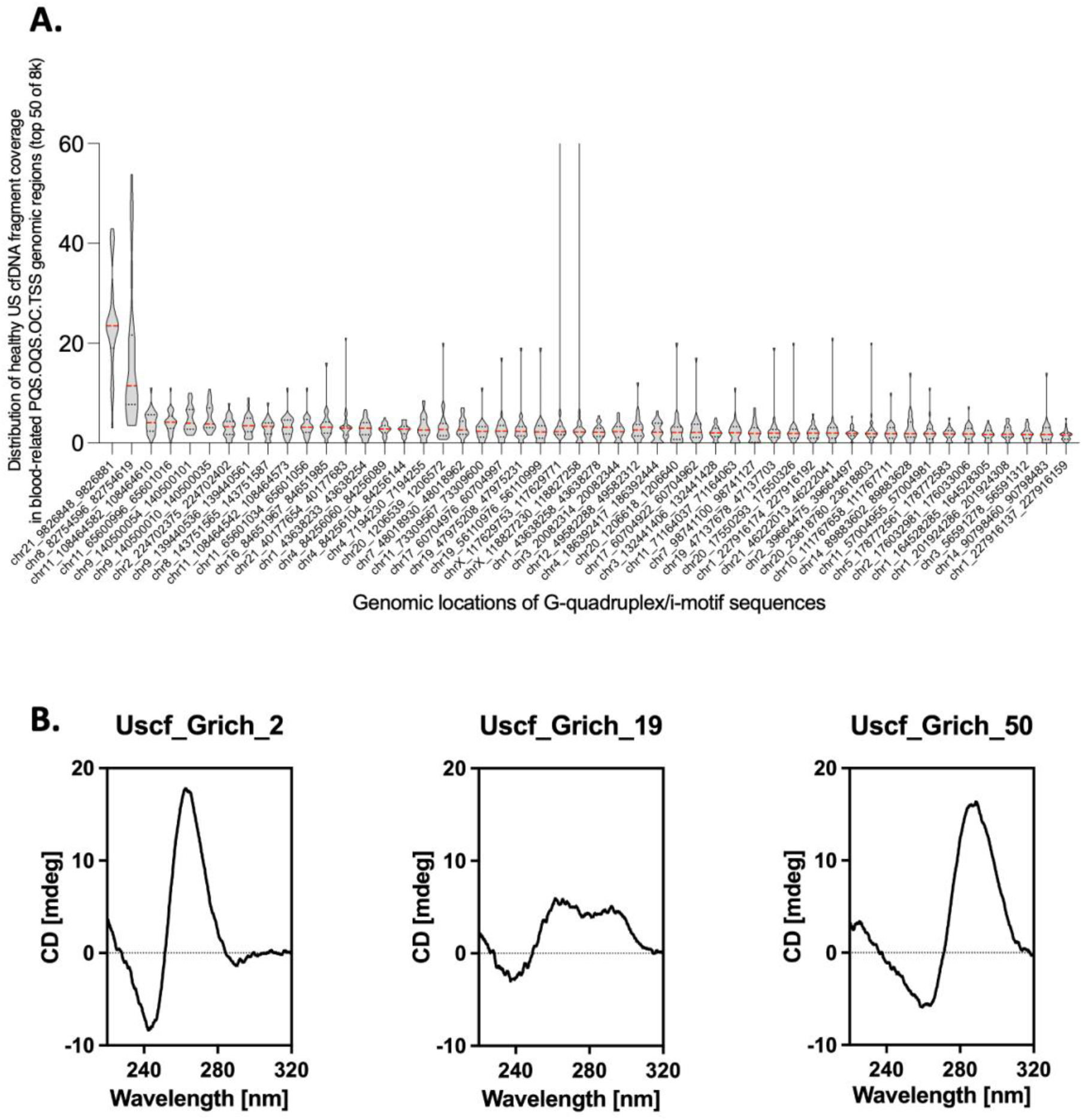
US cfDNA is enriched at putative quadruplex sequences that fold into G4 structures in vitro. A, US cfDNA coverage across blood-related PQS/OQS/OC intervals in plasma from healthy donors. Each violin shows the distribution of CPM-normalized coverage across the 26 sequencing libraries from 20 healthy donors for one interval; intervals are ordered by decreasing median coverage and the 50 highest-ranked of 8,316 qualifying intervals are shown (dashed line, median; dotted lines, quartiles). B, Representative CD spectra of synthetic PQS/OQS/OC sequences, showing spectral signatures consistent with parallel, a mixture of parallel and antiparallel, and antiparallel G4 conformations.

To assess the G4-forming capacity of these 50 highest-ranked regions, we characterized the corresponding synthetic oligonucleotides by circular dichroism (CD) spectroscopy in a buffer mimicking the ionic composition of plasma (cf. Materials and Methods). The CD spectra of the vast majority of the sequences were dominated by a positive band at ~260 nm and a negative band at ~240 nm, consistent with predominantly parallel G4 folding.^15^ A smaller subset showed additional spectral features around ~290 nm, suggesting varying contributions of antiparallel G4 conformations, whereas one sequence displayed a positive band at ~290 nm and a negative band at ~265 nm, characteristic of an antiparallel G4 topology. Representative CD spectra illustrating this range of spectral features are shown in Fig. 1B, with spectra for all 50 sequences provided in Figs. S1 and S2. These measurements demonstrate that sequences representative of the US cfDNA population are intrinsically capable of folding into G-quadruplex structures under plasma-mimicking ionic conditions (132 mM NaCl, 5 mM KCl, 1.5 mM MgCl^2^, 10 mM sodium phosphate, pH 7.4).

Structural competence of isolated synthetic oligonucleotides does not, however, demonstrate that folded G-quadruplexes are present in the nucleic acids that circulate in plasma. We therefore asked whether endogenous plasma nucleic acids adopt G4 structures.

### Folded G4 structures are detected in nucleic acids captured from human blood plasma

To address this, we developed a capture and visualization assay in which no extraction, denaturation or thermal annealing step was applied at any point (Fig. 2A). Terminal deoxynucleotidyl transferase was added directly to plasma, poly(A)-tailing the 3′-hydroxyl termini of endogenous circulating nucleic acids in situ^16^. Tailed plasma was then applied to poly(T)-functionalized magnetic beads or poly(T)-coated microplates, and captured molecules were washed and stained in buffers whose monovalent cation composition remained permissive for G4 folding throughout. This workflow preserves the DNA structural state while providing a universal capture handle through the enzymatically added poly(A) tail, and is closely related to an established strategy for visualizing chromatin features of cfDNA at the single-molecule level, such as nucleosomes carrying specific epigenetic marks^16^. Because circulating nucleic acids were never transferred into a chaotropic or cation-depleted environment, structures present in plasma were not given an opportunity to unfold. Detection under plasma-mimicking and intracellular-mimicking ionic conditions gave equivalent results.

**Fig. 2:**
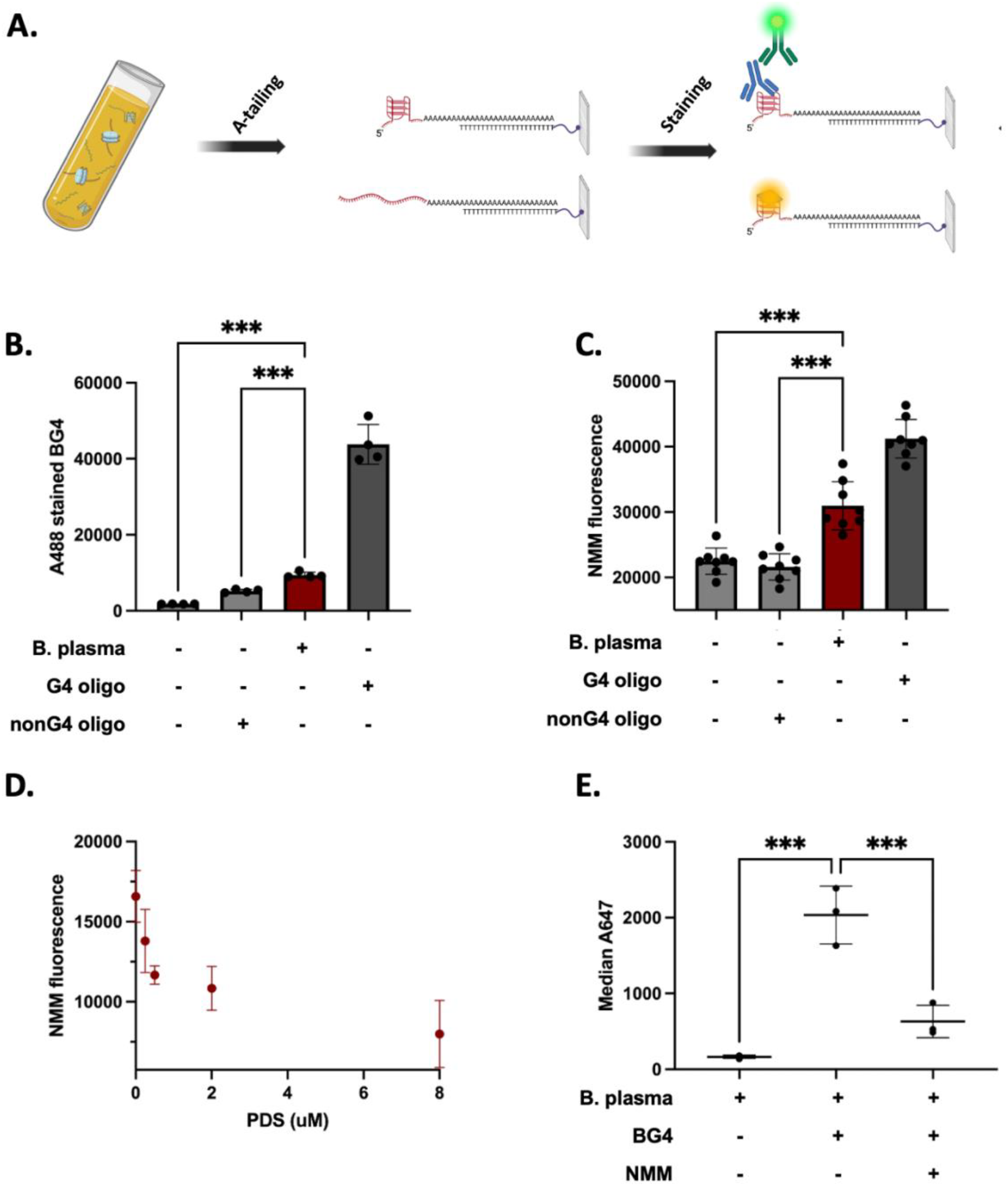
Folded G4 structures are detected in nucleic acids captured from human blood plasma. A, Schematic of the poly(A)-tailing workflow for capturing and staining plasma cfDNA. B, BG4 staining of plasma-captured nucleic acids was significantly higher than that of the non-G4 oligonucleotide and empty controls. The synthetic G4 oligonucleotide served as a positive control for assay performance. Nucleic acids were captured on a poly(T)-coated 96-well plate (n = 4 independent reaction replicates per condition). C, Fluorescence of the G-quadruplex ligand NMM increases significantly in the presence of plasma-captured nucleic acids, relative to the non-G4 oligonucleotide and empty controls. The synthetic G4 oligonucleotide served as a positive control for assay performance. Nucleic acids were captured on a poly(T)-coated 96-well plate (n = 8 independent reaction replicates per condition). For B and C, statistical significance was assessed using Welch’s one-way ANOVA followed by Dunnett’s T3 multiple-comparisons test (P < 0.05). D, NMM specificity confirmed by competition with pyridostatin (PDS), an established small-molecule G4 ligand. Increasing PDS concentrations decrease NMM fluorescence, as expected for competition at G4 binding sites. Plasma cfDNA was captured on poly(T)-coated magnetic beads. (n = 3 independent reaction replicates for 0, 0.25, 0.5, and 8 μM PDS; n = 2 for 2 μM). E, BG4 specificity confirmed by competition with NMM. Plasma cfDNA was captured on poly(T)-coated magnetic beads. BG4 staining increased significantly relative to the no-BG4 control and decreased in the presence of the competing G4 ligand NMM. Statistical significance was assessed using ordinary one-way ANOVA followed by Dunnett’s multiple-comparisons test (P < 0.05; n = 3 independent experimental replicates).

Two oligonucleotide controls were carried through an identical workflow. The positive control was the representative PQS/OQS/OC sequence Uscf_Grich_5 of Fig. 1A (selected for length closely matching the size of US cfDNA), which was captured only after enzymatic A-tailing, confirming that immobilization depends on the added tail rather than on nonspecific surface binding (Fig. 2B). The negative control, derived from a G4 ChIP-qPCR negative-control region in the human ESR1 gene, was A-tailed and captured comparably but cannot fold into a G4. We then probed nucleic acids captured from pooled plasma, pooled to limit donor bias, using BG4, an antibody with well-established specificity for folded G-quadruplex structures^17^. Capture of plasma nucleic acids consistently produced a robust BG4 immunofluorescence signal, whereas controls lacking plasma DNA or BG4 gave significantly lower signals (Fig. 2B). The synthetic G4 control oligonucleotide (Uscf_Grich_5), processed through the complete workflow, gave a reproducibly elevated BG4 signal across replicates, demonstrating that poly(A)-tailing and immobilization are compatible with preservation and detection of pre-existing folded G-quadruplex structures.

To evaluate this signal independently, we used two different G4 ligands, each structurally unrelated to BG4 and to each other. N-methyl mesoporphyrin IX (NMM) is a porphyrin that is weakly emissive in free solution. End-stacking onto a terminal G-tetrad raises its fluorescence quantum yield by orders of magnitude, with pronounced selectivity for parallel over other G4 topologies and especially relative to duplex and single-stranded DNA^18,19^. Pyridostatin (PDS) is a structurally unrelated bis-quinoline ligand that binds the same terminal tetrad surface with high selectivity but is not fluorogenic, which makes the two suitable for reciprocal competition^20,21^. NMM staining of plasma-captured nucleic acids reproduced the BG4 result on both the magnetic-bead and microplate platforms (Fig. 2C, D). Preincubation with PDS reduced NMM fluorescence in a concentration-dependent manner (Fig. 2D), consistent with direct competition between these two G4 ligands. In addition, co-treatment with NMM competed with BG4 detection of plasma-captured fragments, decreasing BG4 fluorescence (Fig. 2E). These mutually consistent competition experiments indicate that recognition depends on specific interaction with folded G-quadruplexes rather than on nonspecific binding to immobilized DNA or to the capture surface.

Two independent capture platforms, two orthogonal detection probes and a third G4 ligand used in competition therefore converge on the same conclusion, providing robust evidence for folded endogenous G-quadruplex structures in human blood plasma. Combined with the computational and structural characterization of US cfDNA-derived sequences, these findings link fragmentomic observations to higher-order DNA structure.

## Discussion

The discovery of US cfDNA introduced a previously unrecognized population of the circulating nucleic acid pool whose molecular organization remains incompletely understood^1–3^. Independent fragmentomic studies associated it with promoter-associated regions enriched in predicted G4-forming sequences, prompting the proposal that higher-order DNA structures contribute to its biology, a model supported until now primarily by fragmentomic observation, genomic localization and computational prediction. By combining structural characterization of US cfDNA-enriched putative quadruplex sequences with direct biochemical interrogation of endogenous plasma nucleic acids, the present study provides experimental evidence for that model.

The central distinction our work draws is between predicted G4-forming potential and experimental structural evidence: sequence composition establishes that a region could fold, not that it does. Synthetic oligonucleotides corresponding to highly ranked US cfDNA-associated PQS regions showed CD signatures consistent with predominantly parallel G4 formation, with a smaller number exhibiting signatures consistent with a mixture of parallel and antiparallel or with antiparallel conformations, supporting their structural competence under plasma-mimicking ionic conditions. Complementing these *in vitro* observations, two orthogonal detection probes, BG4 and NMM, together with reciprocal ligand competition, consistently detected G4-reactive structures in nucleic acids captured from human plasma. These observations extend the interpretation of circulating plasma nucleic acids beyond sequence-derived prediction to biochemical evidence for higher-order nucleic-acid structure.

These findings should also be considered alongside recent reports of extracellular G4 detection. The fluorescent probe G4-Flame suggested the presence of extracellular G-quadruplex structures in human serum, and an increase in G4 levels in cancer patients^22^. Because G4-Flame provides a single fluorescence readout, independent biochemical validation of that serum signal would strengthen its assignment to folded G4 structures.

Very recently, Cabrero-Martín et al. reported elevated G4 signals in plasma from colorectal cancer patients using an electrochemical competitive immunoassay^23^. As reported, the protocol applies a 94 °C denaturation and controlled re-annealing to every sample type, including diluted plasma, and quantifies silica-column extracts. Column-based extraction largely loses the ~50-nucleotide single-stranded cfDNA population, which is recovered only by magnetic-bead extraction^2^. That study therefore reports the G4-forming capacity of plasma-derived DNA under standardized folding conditions, rather than the conformational state of those molecules in blood, a biochemical counterpart to the computational assessment of G4 potential in sequenced cfDNA fragments reported previously^2^. The present work addresses the complementary question: no extraction, denaturation or annealing step is applied at any point, so the structures detected are those already present in plasma.

Here we tested whether G4 structures are present in endogenous, non-denatured nucleic acids captured from human plasma. US cfDNA is, to date, the only single-stranded DNA species reported in human biofluids with G4-forming potential, and several studies have found it depleted of predicted G4-forming sequences in plasma from cancer patients, both across cancer types and within an NSCLC cohort^2,7^. Our experiments do not resolve whether the structures recognized here are formed by DNA or by RNA; the convergence of chemically independent recognition approaches nevertheless supports the presence of folded G-quadruplex structures within this specific circulating population.

Several questions remain beyond the scope of this study. Our findings establish the structural competence of sequences associated with US cfDNA and provide biochemical evidence for G4-reactive structures in plasma-derived nucleic acids, but they do not determine the biological origin of these structures, whether G4 formation precedes DNA fragmentation, or whether folding contributes to fragment stability after release into the circulation. Nor do we propose that all US cfDNA adopts G4 conformations. Rather, our results establish that folded G4 structures are present within this molecular population, and provide an experimental framework for investigating their abundance, biological function, and clinical relevance.

In summary, this work provides direct biochemical evidence for folded G4 structures in nucleic acids circulating in human blood plasma and shows that structural hypotheses arising from fragmentomic analysis can be tested experimentally by combining computational prediction, biophysics and biochemistry with chemically orthogonal probes for molecular recognition. These findings support the view that higher-order nucleic-acid structures may represent an additional molecular dimension of circulating nucleic acids, beyond their sequence and fragment-size characteristics. Extending fragmentomics from sequence-based observation toward direct structural interrogation is an important step toward understanding the higher-order molecular architecture of circulating nucleic acids.

## Materials and Methods

### Reagents and Materials

Terminal deoxynucleotidyl transferase (Terminal Transferase, Cat. No. M0315L), TdT reaction buffer, CoCl_2_ solution and Oligo d(T)^25^ Magnetic Beads (Cat. No. S1419S) were purchased from New England Biolabs (Ipswich, MA, USA). dATP (100 mM stock solution; Cat. No. U1201) was obtained from Promega (Madison, WI, USA). Pierce™ NeutrAvidin™ Coated Plates (Cat. No. 15117) and SuperBlock™ Blocking Buffer in PBS (Cat. No. 37580) were purchased from Thermo Fisher Scientific (Waltham, MA, USA). Dulbecco’s phosphate-buffered saline (DPBS) without Ca^2+^ and Mg^2+^ (Cat. No. D8537), Tween® 20 (Cat. No. P2287), and salmon sperm DNA (Cat. No. 15632011) were purchased from Sigma-Aldrich (St. Louis, MO, USA). N-Methyl Mesoporphyrin IX (NMM; Cat. No. 2P5Y.1) was obtained from Carl Roth (Karlsruhe, Germany). Pyridostatin hydrochloride (PDS; Cat. No. SML2690; ≥98% HPLC) was purchased from Sigma-Aldrich (St. Louis, MO, USA). DYKDDDDK Tag Antibody (anti-FLAG; Cat. No. 2368) was purchased from Cell Signaling Technology (Danvers, MA, USA), and Alexa Fluor™ 488 goat anti-rabbit IgG (H+L) (Cat. No. A11008) and Alexa Fluor™ 647 goat anti-rabbit IgG (H+L) (Cat. No. A21244) were obtained from Invitrogen (Thermo Fisher Scientific).

Unless otherwise stated, PBST refers to PBS supplemented with 0.05% (v/v) Tween-20, and poly(T)^25^ refers to a 25-mer oligo-deoxythymidine.

### Human Plasma

A pooled human plasma from healthy female donors, filtered through a 0.2 μm membrane (BioIVT, Cat. No. HUMANPLK2-8905-D), was used throughout the study. The plasma samples were stored at −80 °C, thawed at 4 °C and centrifuged at 14,000 × *g* for 10 min at 4 °C to minimize the contamination of cellular DNA.

### Computational Enrichment Analysis of US cfDNA at Putative Quadruplex Sequences

Putative quadruplex sequences (PQS) were predicted genome-wide in hg19 using fastaRegexFinder.py with the parameters --quiet -r ‘([gG]{3,}\w{1,7}){3,}[gG]{3,}’, corresponding to four or more G-runs of at least three guanines separated by loops of 1–7 nucleotides. Experimentally observed quadruplex sequences (OQS) were obtained from G4-seq^10^ (Gene Expression Omnibus accession GSE63874); the four available hit files were concatenated and unified using the BEDTools functions mergeBed and sortBed. PQS overlapping an OQS (PQS/OQS) were retrieved with bedtools intersect (-u). Blood-related open chromatin was retrieved from ChIP-Atlas^24^ (https://chip-atlas.org) using antigen class DNase-seq, cell-type class blood, significance threshold 500 and all blood cell types, then sorted and merged. Transcription start sites (TSS) were annotated from the hg19 gene annotation as described previously^11^, taking the most 5′ transcript start per gene on each strand. Each TSS was extended by 1 kb in both directions and overlapping windows belonging to the same gene were merged to generate hg19 promoter intervals. Successive intersections of PQS/OQS with blood-related open chromatin and then with promoter intervals yielded 8,316 PQS/OQS/OC intervals (median width 25 bp, range 15–175 bp).

Ultrashort cfDNA coverage was computed from the healthy-donor magnetic-bead single-stranded DNA (MB-ssDNA) libraries reported by Hudecova et al. (2022)^2^, available from the European Genome-Phenome Archive under accession EGAS00001005093. Paired-end aligned reads were size-selected in silico to fragments of 30–70 bp, duplicate-marked and blacklist-filtered, and converted into CPM-normalized coverage tracks using deepTools bamCoverage (--normalizeUsing CPM). Coverage of each of the 26 MB-ssDNA libraries, derived from 20 healthy donors (16 donors contributed a single library and four contributed two or three replicate libraries from the same plasma sample; all samples were from a single timepoint), was then quantified across the 8,316 PQS/OQS/OC intervals using deepTools multiBigwigSummary in BED-file mode (--outRawCounts), yielding a region × library matrix of CPM values. Intervals were ranked by their median CPM coverage across the 26 libraries, and the 50 highest-ranked intervals are shown in Fig. 1A as violin plots of the per-library CPM distribution for each interval (dashed line, median; dotted lines, quartiles). Genomic sequences of the highest-ranked intervals were extracted against hg19 with bedtools getfasta and used to design the synthetic oligonucleotides listed below. Interval operations were performed with BEDTools^25^ and coverage operations with deepTools^26^.

### DNA Oligonucleotides

#### Oligonucleotides for CD screening

DNA oligonucleotides were purchased from Generi Biotech (Czech Republic). Representative G-rich sequences were selected from the highly abundant ultrashort plasma cfDNA fragments reported by Hudecova *et al*. (Fig. 1A) as described above. The trinucleotides ATT and TTA were added to the 5′ and 3′ ends, respectively, to minimize intermolecular G4 formation.

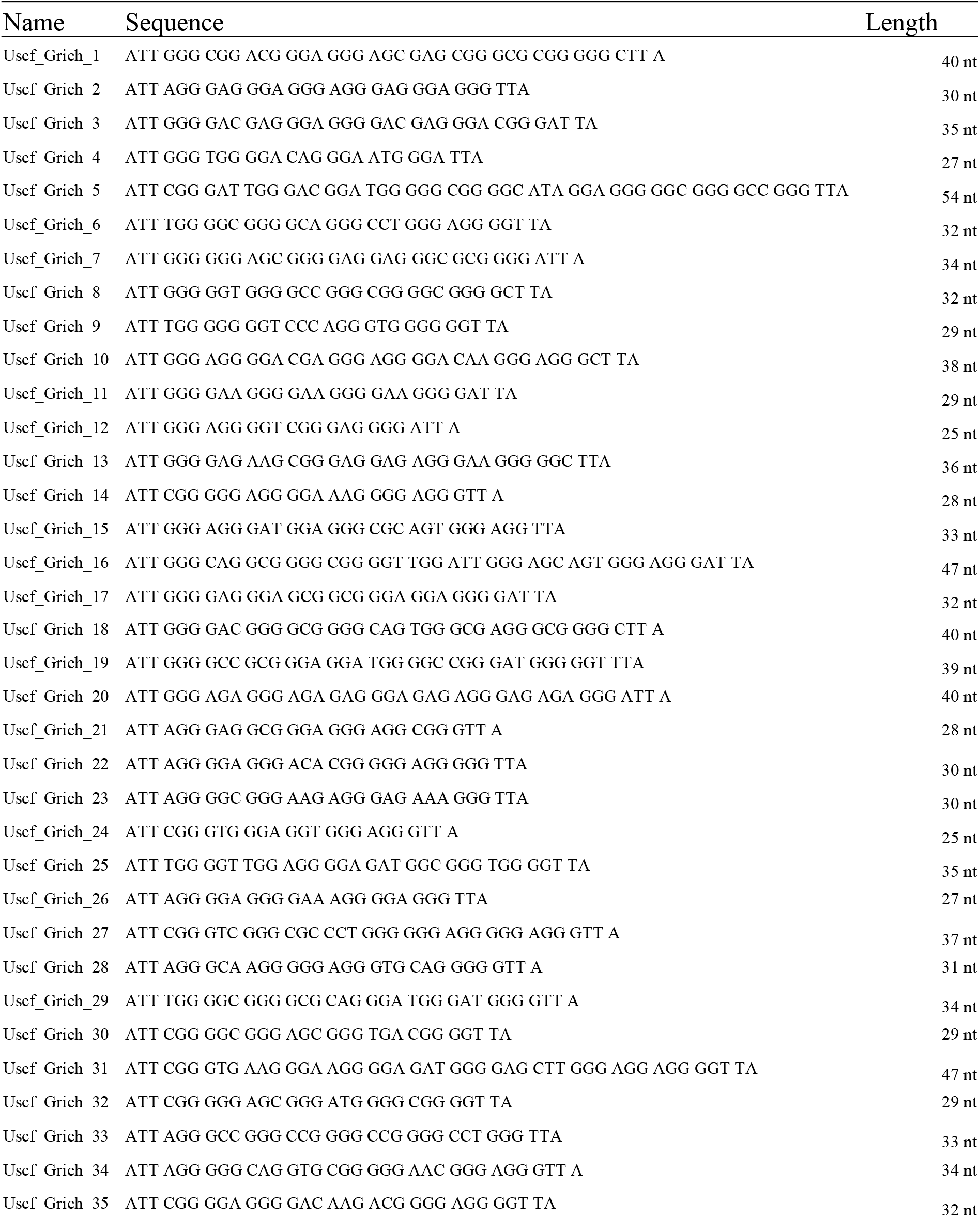
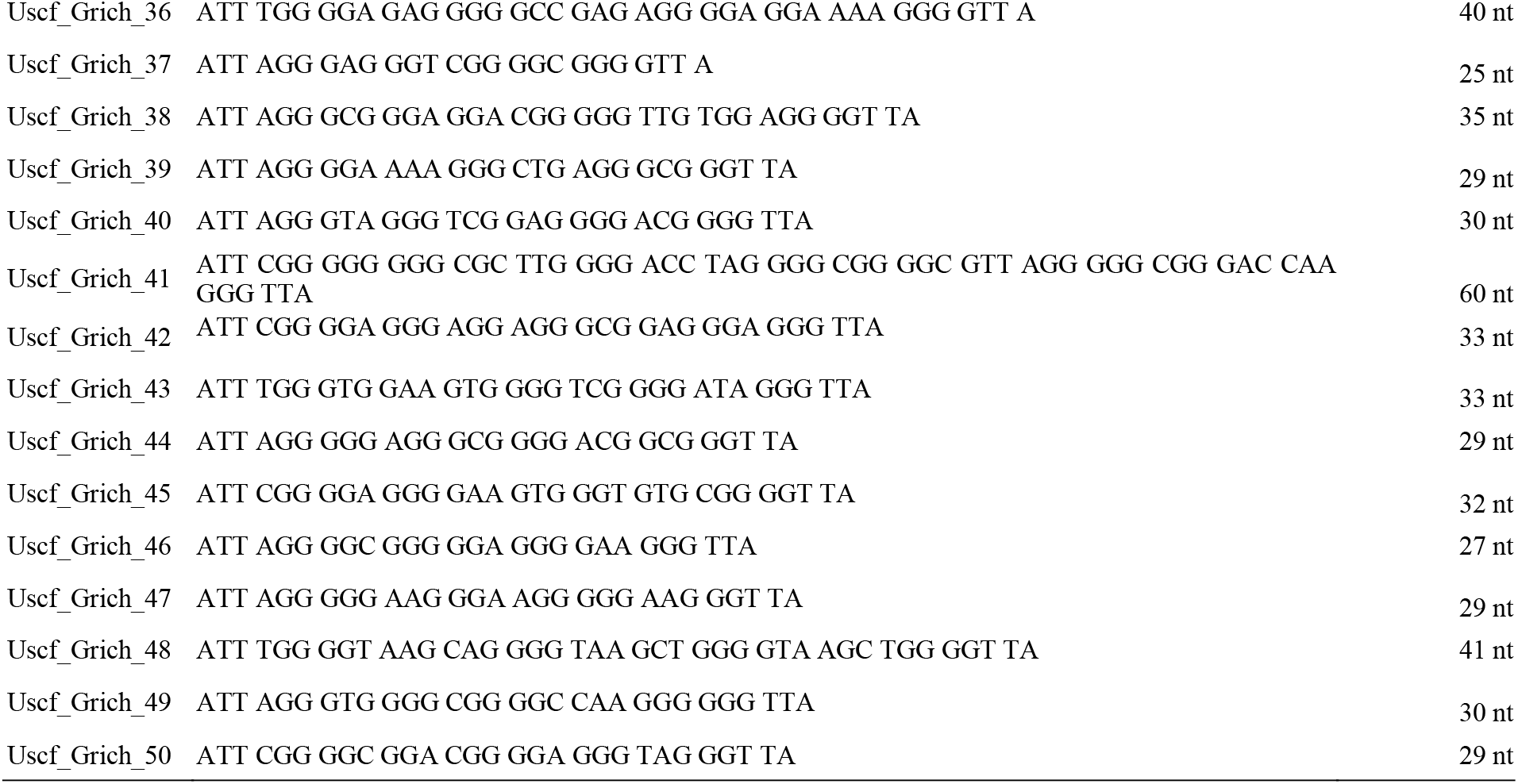

#### Oligonucleotides for detection assays

Synthetic DNA oligonucleotides were purchased from Integrated DNA Technologies (IDT, Leuven, Belgium) with standard purification and supplied at 100 μM in TE buffer.

A representative G-rich sequence was selected from the highly abundant ultrashort plasma cfDNA fragments reported by Hudecova et al. A non-G4-forming sequence of comparable length served as the negative control, and a 5′-biotinylated poly(T)^25^ oligonucleotide was used to generate poly(T)-functionalized capture surfaces.

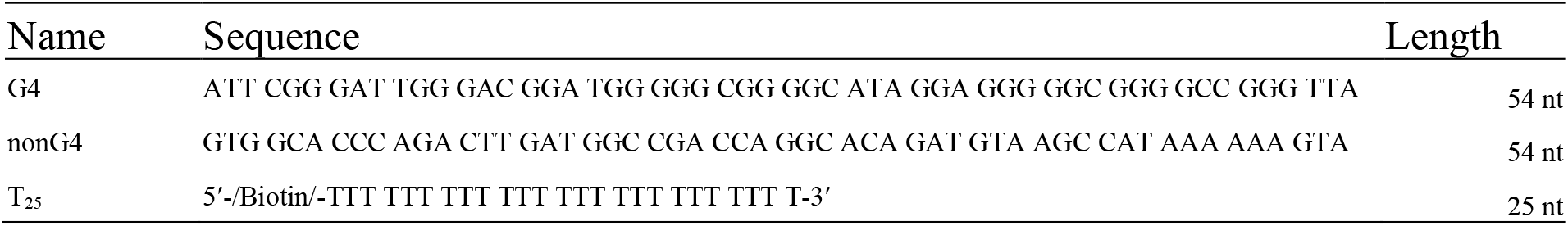

### Folding of Synthetic DNA Oligonucleotides

Synthetic oligonucleotides were diluted to 0.1 μM in phosphate-buffered saline (PBS), heated to 95 °C for 10 min, and allowed to cool slowly to room temperature overnight to permit equilibrium folding. Folded oligonucleotides were used for biophysical characterization and as positive controls throughout the biochemical experiments.

### Circular Dichroism Spectroscopy

Circular dichroism experiments were performed on a JASCO J-815 spectropolarimeter equipped with a JASCO PTC-423S temperature controller using a 0.1 cm path-length quartz cuvette. The oligonucleotides were dissolved in water to form 1 mM stock solutions. Then 20 µM oligonucleotide solution was prepared for each sample in blood plasma mimicking buffer (132 mM NaCl, 5 mM KCl, 1.5 mM MgCl^2^, 10 mM sodium phosphate, pH 7.4). Samples were heated at 95 °C for 5 min and let cool down overnight. CD spectra were recorded at 25 °C in a wavelength range of 220–320 nm using the following parameters: scanning speed of 100 nm/min; bandwidth of 2 nm; data interval of 0.1 nm; response of 1 s. The buffer contribution was subtracted from each CD spectrum after the acquisition. Raw experimental data were processed using JASCO Spectra Manager 2.09.10 software and then analyzed using GraphPad Prism 8.

### Preparation of Poly(T)-Functionalized Capture Surfaces

#### Microplate Capture Format

Pierce™ NeutrAvidin™ Coated Plates were washed twice with 200 μL PBST per well. Biotinylated poly(T)^25^ oligonucleotides were diluted to 6 μM in PBST supplemented with 10% (v/v) SuperBlock™ Blocking Buffer, and 105 μL was added to each well. Plates were sealed and incubated overnight at room temperature on an orbital shaker (120 rpm) to allow immobilization of the poly(T)^25^ capture oligonucleotide. Prior to use, wells were washed three times with 200 μL PBST.

#### Magnetic Bead Capture Format

Oligo d(T)^25^ Magnetic Beads were prepared according to the manufacturer’s instructions before hybridization with poly(A)-tailed DNA.

### Terminal Transferase-Mediated Poly(A)-Tailing

Native plasma cfDNA and synthetic oligonucleotides were poly(A)-tailed using terminal deoxynucleotidyl transferase (TdT).

Each 100 μL reaction contained 20 μL plasma (or synthetic oligonucleotide solution), 1× TdT reaction buffer, 2.5 mM CoCl^2^, 42 μM dATP and 2 μL TdT. Reactions were incubated for 1 h at 37 °C and terminated with EDTA at a final concentration of 30 mM.

No denaturing conditions were introduced at any point following A-tailing, or during subsequent capture and staining.

### Native Plasma DNA Capture

Immediately after A-tailing, DNA was captured by hybridization to immobilized poly(T)^25^.

For the microplate format, A-tailed samples were transferred directly to poly(T)-functionalized wells and incubated for 1 h at room temperature on an orbital shaker (120 rpm).

For the magnetic bead format, A-tailed DNA was incubated with 0.5 µL Oligo d(T)^25^ Magnetic Beads for 1 h at room temperature on a rotating wheel.

The capture workflow was adapted from the native plasma DNA immobilization strategy described previously^16^, replacing the original imaging surface with poly(T)-functionalized microplates or magnetic beads for downstream biochemical interrogation.

### BG4 Immunodetection

Captured DNA was incubated with recombinant BG4 antibody (final concentration 0.03 µM), produced in-house according to the original protocol of Biffi *et al*. ^17^, in PBST supplemented with 10% SuperBlock™ Blocking Buffer and salmon sperm DNA (1:2000).

Salmon sperm DNA was included as a nonspecific competitor DNA to minimize background interactions of BG4 with immobilized DNA while preserving selective recognition of folded G-quadruplex structures.

BG4 incubation was performed for 45 min at room temperature. Microplate assays were incubated on an orbital shaker (120 rpm), whereas magnetic bead assays were incubated on a rotating wheel. Samples were subsequently washed three times with PBST.

Bound BG4 was detected using anti-FLAG antibody (1:400) followed by Alexa Fluor™ 488 goat anti-rabbit IgG (1:1000). Both antibody incubations were performed for 30 min at room temperature using the same incubation format (orbital shaker for microplates or rotating wheel for magnetic beads), with three PBST washes between each step.

Alexa Fluor™ 488 fluorescence was measured using a Tecan Spark multimode microplate reader (Tecan, Switzerland) in top-read fluorescence mode with 485 nm excitation (20 nm bandwidth) and 535 nm emission (20 nm bandwidth). Measurements were acquired using an automatic dichroic 510 nm mirror.

For statistical analysis of the microplate-based BG4 assay, four independently prepared reaction wells were analyzed for each condition (n = 4 experimental replicates). The Empty and non-G4 oligonucleotide conditions served as negative controls, whereas the G4 oligonucleotide was included as a positive control to verify assay performance and was excluded from statistical hypothesis testing. Differences among the Empty, non-G4 oligonucleotide, and blood plasma conditions were assessed using Welch’s one-way ANOVA, followed by Dunnett’s T3 multiple-comparisons test. Statistical significance was defined as *P < 0*.*05*. Data were analyzed using GraphPad Prism 10.

### N-Methyl Mesoporphyrin IX Detection

Captured DNA was incubated with N-methyl mesoporphyrin IX (NMM) at a final concentration of 0.1 μM for 30 min at room temperature. Microplate assays were incubated on an orbital shaker, magnetic bead assays on a rotating wheel.

NMM fluorescence was measured using a Tecan Spark multimode microplate reader (Tecan, Switzerland) in top-read fluorescence mode with 399 nm excitation (10 nm bandwidth) and 610 nm emission (10 nm bandwidth). Measurements were acquired using an automatic dichroic 510 nm mirror, 30 flashes per well, and an integration time of 40 µs.

For the microplate-based NMM assay, eight independently prepared reaction wells were analyzed for each experimental condition (n = 8). The Empty and non-G4 oligonucleotide conditions served as negative controls, whereas the G4 oligonucleotide was included as a positive control to verify assay performance and was excluded from statistical hypothesis testing. Differences among the Empty, non-G4 oligonucleotide, and blood plasma conditions were assessed using Welch’s one-way ANOVA, followed by Dunnett’s T3 multiple-comparisons test. Statistical significance was defined as *P < 0*.*05*. Data were analyzed using GraphPad Prism 10.

### Pyridostatin Competition Assay

To evaluate the specificity of NMM binding, plasma-derived poly(A)-tailed DNA was first captured on Oligo d(T)25 Magnetic Beads as described above. Following capture and washing, bead suspensions were aliquoted into individual wells and incubated for 1 h at room temperature on an orbital shaker (120 rpm) with increasing concentrations of pyridostatin (PDS; 0, 0.25, 0.5, 2.0 or 8.0 μM). NMM was then added to a final concentration of 1 μM and fluorescence was measured on a HIDEX Sense multimode microplate reader in top-read mode (390/20 nm excitation, 610/20 nm emission; 10 excitation flashes per well, 50% lamp power). PDS competition data were plotted as mean fluorescence signal ± standard deviation (SD). The number of independent reaction replicates was n = 3 for 0, 0.25, 0.5, and 8 µM PDS and n = 2 for 2 µM PDS. Graphs were generated using GraphPad Prism 10.

### BG4 Competition Assay

For competition experiments, plasma-derived poly(A)-tailed DNA was captured on Oligo d(T)^25^ Magnetic Beads as described above. Following capture and washing, bead-bound DNA was pre-incubated with 10 μM N-Methyl Mesoporphyrin IX (NMM) for 30 min at room temperature on a rotating wheel in PBS buffer. BG4 antibody was subsequently added directly to the reaction (0.012 µM) without removal of the ligand, and both components were co-incubated for 1 h on a rotating wheel. Beads were washed twice with PBS buffer before sequential staining with anti-FLAG antibody (1:800 in PBS buffer) and Alexa Fluor™ 647-conjugated goat anti-rabbit IgG (1:1000 in PBS buffer), each followed by two washes with PBS buffer. Finally, beads were resuspended in 400 μL FACS buffer (DPBS supplemented with 2 mM EDTA) prior to flow cytometric analysis.

Fluorescence intensities of stained magnetic beads were measured using a BD FACSCanto II flow cytometer and analyzed using FlowJo™ v11 (BD Life Sciences). For statistical analysis, the median Alexa Fluor 647-A fluorescence intensity was determined for each independent experimental replicate. Differences among the three conditions (no BG4, BG4, and NMM– BG4) were assessed using one-way ANOVA followed by Dunnett’s multiple-comparisons test, with the BG4 condition used as the reference. Statistical significance was defined as *P* < 0.05. Data were analyzed using GraphPad Prism 10.

## Acknowledgements

We thank the ITCC (IT Center University of Cologne, formerly RRZK) for providing compute resources on the DFG-funded HPC (High Performance Computing) system RAMSES (Research Accelerator for Modeling and Simulation with Enhanced Security) as well as support (DFG funding number: INST 216/512-1 FUGG).

We thank Michala BuČková for assistance with the CD measurements. We acknowledge CIISB, Instruct-CZ Centre, supported by MEYS CR (LM2023042) and by the European Regional Development Fund project “Innovation of Czech Infrastructure for Integrative Structural Biology” (No. CZ.02.01.01/00/23_015/0008175).

We thank the Center for Molecular Medicine Cologne (CMMC) flow cytometry facility for access to the FACS Canto II.

Ultrashort cfDNA sequencing data were generated in the study of Hudecova et al. (2022) and are available from the European Genome-Phenome Archive under accession EGAS00001005093.

## Author contributions

Conceptualization: R.H.-H.

Methodology: M.G., O.V.R., T.A., P.H., A.C., J.-L.M., L.T., R.H.-H.

Investigation: M.G., O.V.R., A.C.

Visualization: M.G., O.V.R., A.C.

Funding acquisition: R.H.-H., L.T., J.-L.M.

Project administration: R.H.-H.

Supervision: R.H.-H., L.T., J.-L.M.

Writing – original draft: M.G., R.H.-H.

Writing – review & editing: M.G., O.V.R., T.A., P.H., A.C., J.-L.M., L.T., R.H.-H.

---

**Fig. S1:**
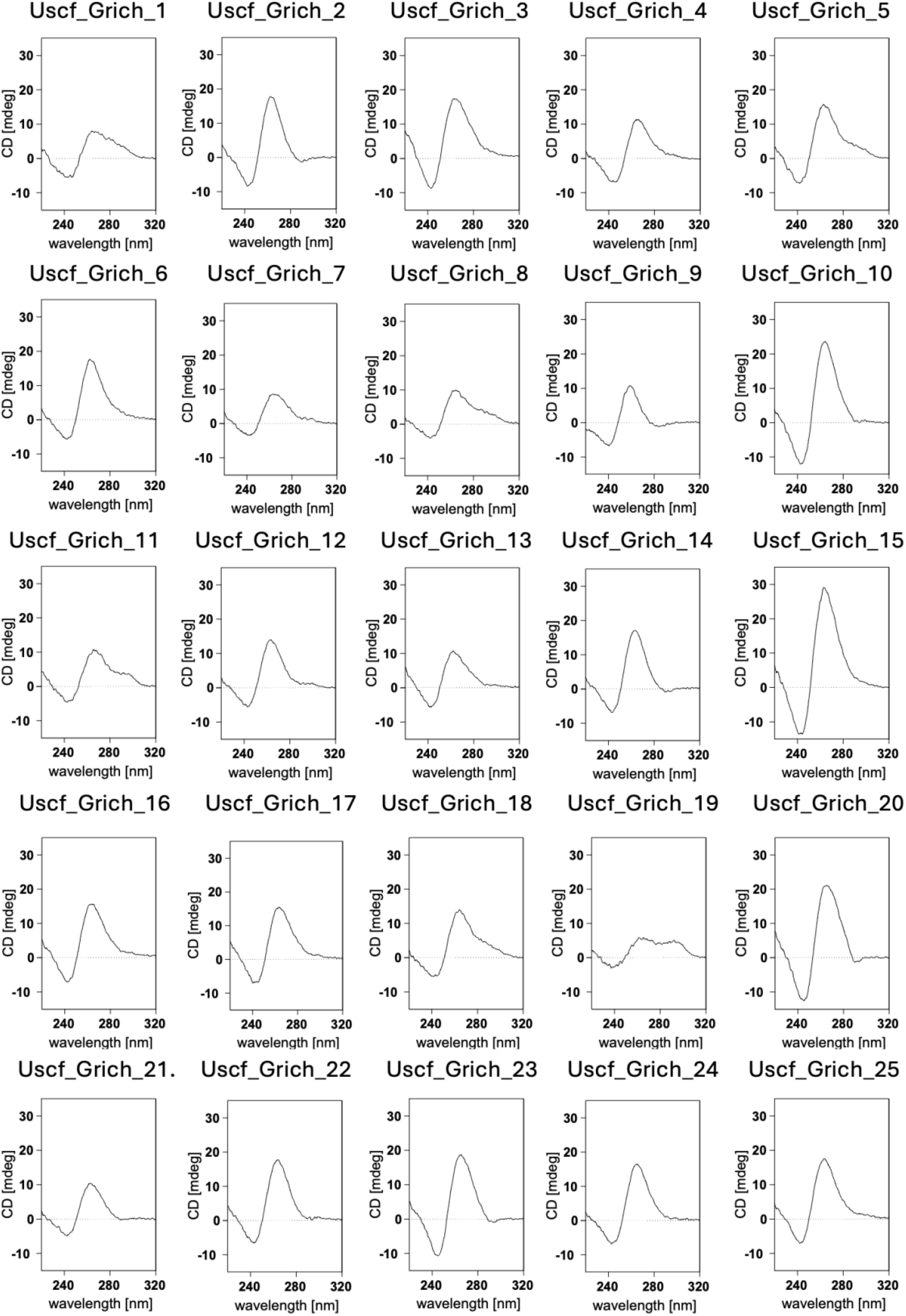
CD spectra acquired at 25 °C for Uscf_Grich1–25 (cf. Materials and Methods) in a blood plasma-mimicking buffer (132 mM NaCl, 5 mM KCl, 1.5 mM MgCl_2_, 10 mM sodium phosphate, pH 7.4) at an oligonucleotide concentration of 20 µM.

**Fig. S2:**
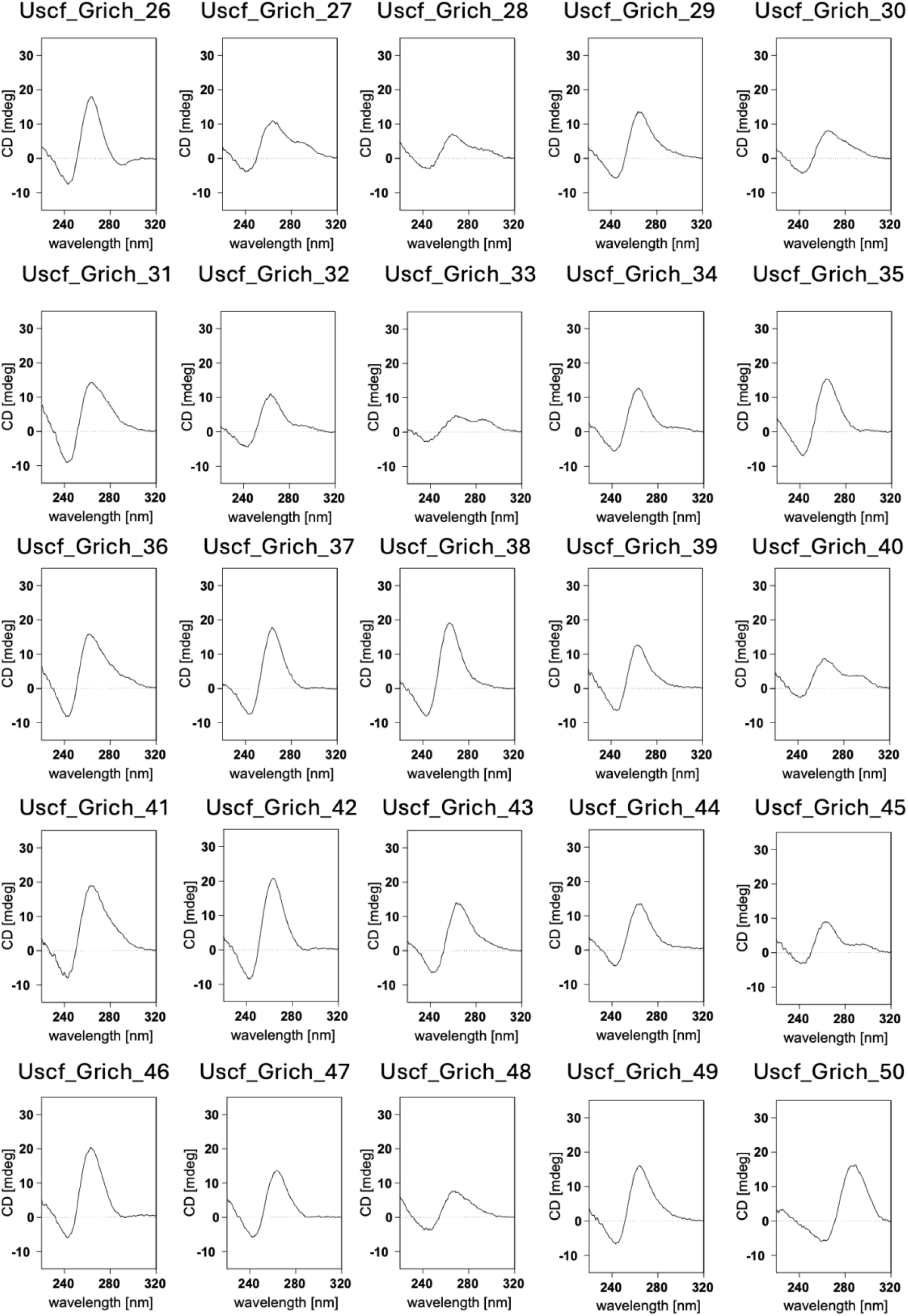
CD spectra acquired at 25 °C for Uscf_Grich26–50 (cf. Materials and Methods) in a blood plasma-mimicking buffer (132 mM NaCl, 5 mM KCl, 1.5 mM MgCl_2_, 10 mM sodium phosphate, pH 7.4) at an oligonucleotide concentration of 20 µM.

